# Measurement Error Correction of Genome-Wide Polygenic Scores in Prediction Samples

**DOI:** 10.1101/165472

**Authors:** Elliot M. Tucker-Drob

## Abstract

DiPrete, Burik, & Koellinger (2017; http://dx.doi.org/10.1101/134197) propose using an instrumental variable (IV) framework to correct genome-wide polygenic scores (GPSs) for error, thereby producing disattenuated estimates of SNP heritability in predictions samples. They demonstrate their approach by producing two independent GPSs for Educational Attainment (“multiple indicators”) in a prediction sample (Health and Retirement Study; HRS) from independent sets of SNP regression weights, each computed from a different half of the discovery sample (EA2; Okbay et al. 2016), i.e. “by randomly splitting the GWAS sample that was used for [the GPS] construction.”

Here, I elucidate how a structural equation modeling (SEM) framework that specifies true score variance in GPSs as a latent variable can be used to derive an equivalent correction to the IV approach proposed by DiPrete et al. (2017). This approach, which is rooted in a psychometric modeling tradition, has a number of advantages: (1) it formalizes the assumed data-generating model, (2) it estimates all parameters of interest in a single step, (3) is can be flexibly incorporated into a larger multivariate analysis (such as the “Genetic Instrumental Variable” approach proposed by DiPrete et al., 2017), (4) it can easily be adapted to relax assumptions (e.g. that the GPS indicators equally represent the true genetic factor score), and (5) it can easily be extended to include more than two GPS indicators. After describing how the multiple indicator approach to GPS correction can specified as a structural equation model, I demonstrate how a structural equation modeling approach can be used to correct GPSs for error using SNP heritability obtained using GREML or LD score regression to produce a correction that is equivalent to an approach recently proposed by Daniel Benjamin and colleagues. Finally, I briefly discuss what I view as some conceptual limitations surrounding the error correction approaches described, regardless of the estimation method implemented.

## Instrumental Variable Analysis for Causal Inference

For the reader unfamiliar with IV analysis, I begin with a brief overview of the basics of the IV analysis. Under an IV approach, variation in an instrument induces variation in a factor that is hypothesized to have a causal effect on an outcome of interest. A key assumption (termed the *exclusion* restriction) is that, beyond the mediating role of the hypothesized causal factor, the instrument is uncorrelated with the outcome. If this assumption is correct, then an IV approach can be leveraged to produce a consistent estimate the casual effect of the hypothesized causal factor on the outcome.

An example of IV analysis is a Raising of the School Leaving Age (ROSLA) or minimum compulsory schooling policy that is implemented (or rolled out at different times) pseudo randomly across municipalities (e.g. Brinch and Galloway, 2012). The key identifying assumption (the exclusion restriction) is that the policy is not directly correlated with IQ above and beyond its effect via years of education (EduYears). A traditional two stage least squares (2SLS) approach to estimating the causal effect of EduYears on IQ involves saving the predicted values from a regression of EduYears on regional and/or temporal variation in the ROSLA policy (Stage 1). These values are then used to predict IQ (Stage 2). Covariates (e.g. time) should be included in both stages if the exclusion restriction is only plausible conditional on them. The regression coefficient estimated in the second stage represents the local average treatment effect of a year of education on IQ.

A less commonly used method of estimating an IV analysis is in a single step as a structural equation model. Figure 1 contains a path diagram representation of this structural equation model using Reticular Action Model (RAM) notation (Boker, McArdle, & Neale, 2002). This model makes the exclusion restriction explicit, as ROSLA is not allowed to correlate with the IQ residual. Note that structural equation models can be estimated using several different methods (Muthén & Muthén, 2017), perhaps the most popular of which is maximum likelihood. For instance, bootstrapping may be used when there is concern that parameters may have non-normal sampling distributions.

**Figure 1.**
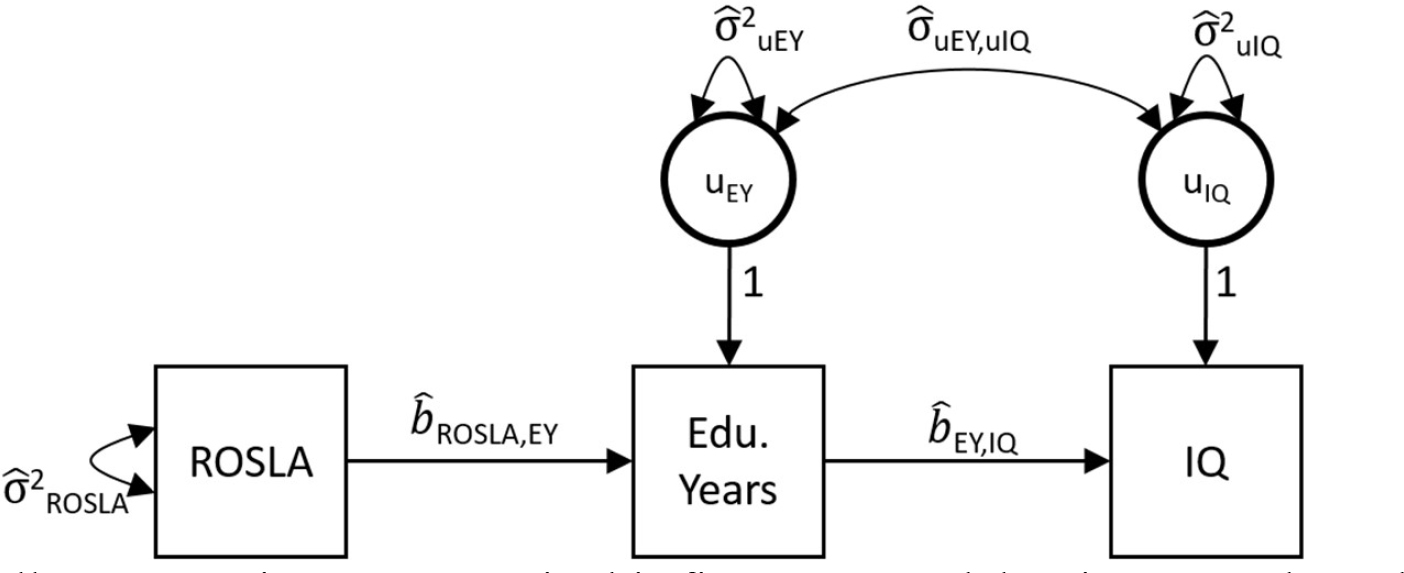
Note that all nonnumeric parameters in this figure are model estimates and are denoted by a hat (^). Numeric parameters are fixed values. Basic statistics that are calculated directly from the sample data have no hats.

The regression structure of this model, excluding means and intercepts, can be written as:

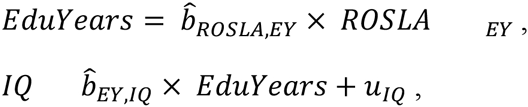

And the covariance structure of the independent variables in this model can be written as:

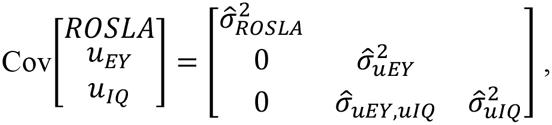

where model-estimated parameters are denoted by a hat (^), and basic statistics calculated directly from the sample have no hats. Variances are denoted by σ, covariances are denoted by σ^2^, and regression coefficients are denoted by *b*.

This model implies that

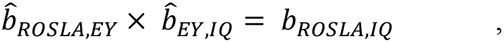

and

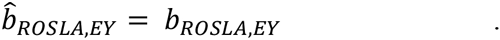

Therefore the key parameter 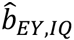 can be calculated as the ratio of two univariate regression coefficients from the sample data, which is equivalent to the ratio of the two corresponding covariances:

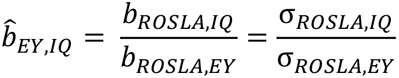

Under this approach, if the ROSLA policy raises average EduYears by .5 years and the average IQ by 1.5 points, then the local average treatment effect of 1 year of education is a raise in IQ of 3 points (1.5/.5 = 3). Thus, the IV approach can be conceptualized as a form of rescaling the “treatment effect” of the instrument on the dependent variable (IQ) in units of the explanatory factor (EduYears). The 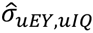 parameter represents unmeasured confounding between EduYear and IQ, which is independent of the instrument and causally ambiguous.

We can arrive at this same result using the 2SLS, in which the regressions are estimated separately. The expected value of EduYears from its regression on ROSLA (so called “Stage 1”) is substituted into the regression Equation for IQ (so called “Stage 2”) yielding what is often referred to as the reduced form equation:

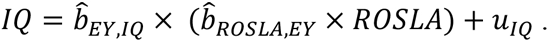

Given that the regressions of IQ on ROSLA and EduYears on ROSLA can be directly computed from the sample data as

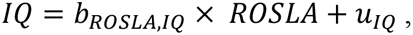

and

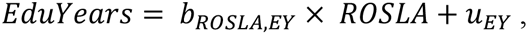

it follows that

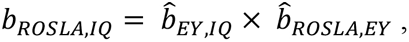

such that

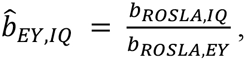

just as is implied by the structural equation model. Moving forward, I use the structural equation modeling framework, rather than the 2SLS framework.

## Instrumental Variable Analysis for Error Correction

DiPrete et al. (2017) note that when multiple indicators with independent sources of error are available, these indicators can be used as instruments for one another other to correct for measurement error. Figure 2 illustrates how such an approach works in the context of the association between educational attainment (EduYears) and IQ, with EduYears measured imperfectly by two independent measures (e.g. mother and father reports of offspring EduYears).

**Figure 2.**
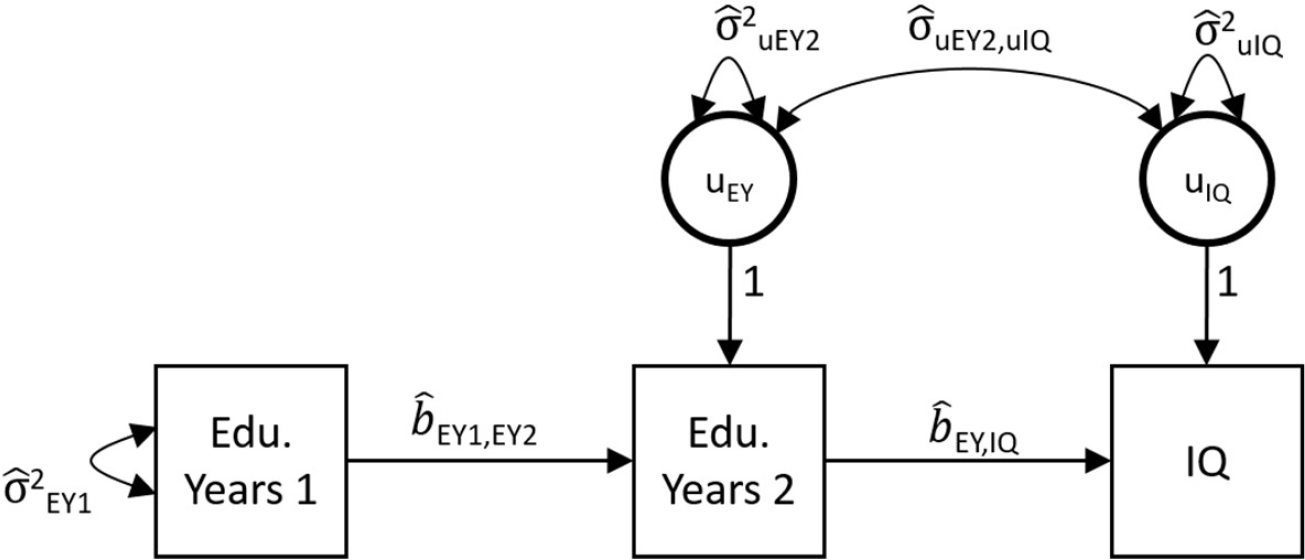
Note that all nonnumeric parameters in this figure are model estimates and are denoted by a hat (^). Numeric parameters are fixed values. Basic statistics that are calculated directly from the sample data have no hats.

In Figure 2, EduYears1 is used as an instrument for EduYears2 to remove measurement error from the unstandardized estimate of the EduYears➔IQ association. The regression coefficient b_EY,IQ_ represents the causal effect of one additional year of education on IQ in IQ points per year. The justification for the exclusion restriction is that EduYears1 is only independent of EduYears2 because of random measurement error. Therefore, EduYears1 is not expected to be associated with the IQ independent of the disattenuated effect of EduYears2.

Interestingly, in this model, the term σuEY2,uIQ is expected to be negative, as it represents the downward bias (attenuation) of the association between EduYears2 and IQ that would have resulted from measurement error had it not been corrected for.

Under this model:

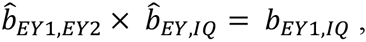

and

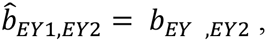

Or alternatively put

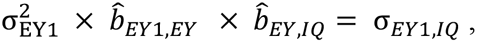

which reduces to

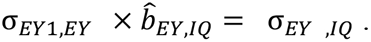

Therefore,

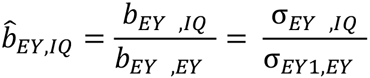

For example, if imperfectly measured EduYears is associated with IQ at 1 point per year and imperfectly measured EduYears predicts another imperfect measure of Eduyears at .8 points/year, then 1 year of education raises IQ by 1.25 points (1/.8 = 1.25). Importantly, in this multiple indicator context, the 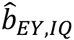 term is corrected for unreliability, but does not necessarily represent a causal effect of EduYears on IQ.

## An Explicit Psychometric Model for Error Correction

Figure 3 represents an alternative modelling strategy for the same multiple indicator data. As was the case earlier, EduYears1 and EduYears2 are two imperfect measures of EduYears (e.g. mother and father reports of offspring EduYears). We create a latent variable representing true score variance in EduYears, anchoring the metric of the variable to the metric of manifest variable EduYears1 via a fixed unstandardized loading of 1. As EduYears1 and EduYears2 are expected to contain equal amounts of true score variance, we make the simplifying (but unnecessary) assumption that EduYears1 and EduYears2 load equally on the latent variable. Under this model,

**Figure 3.**
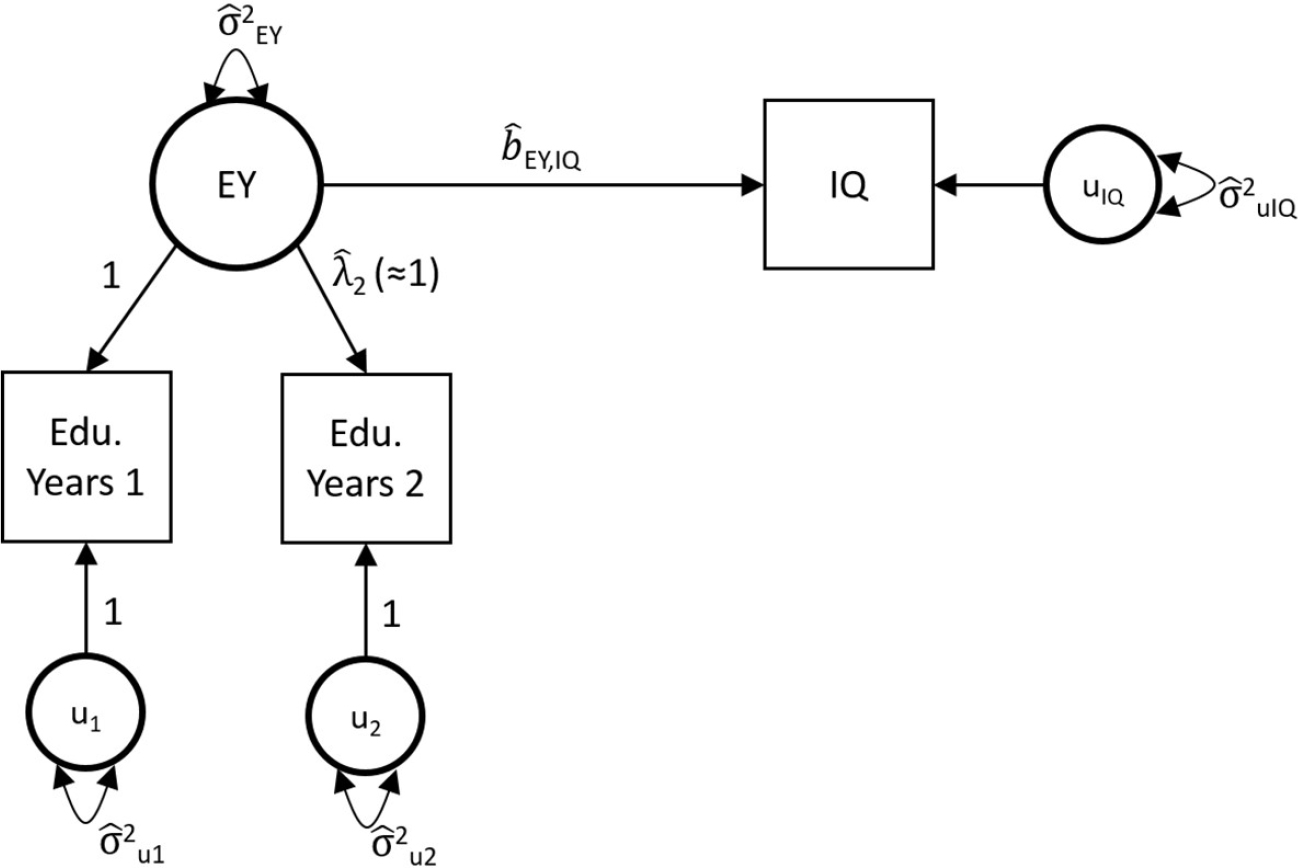
Note that all nonnumeric parameters in this figure are model estimates and are denoted by a hat (^). Numeric parameters are fixed values. Basic statistics that are calculated directly from the sample data have no hats.

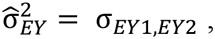

and

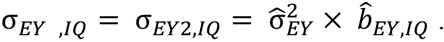

Thus

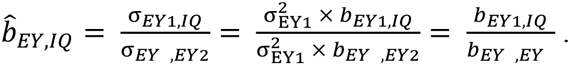

This estimate of b_EY,IQ_ is exactly the same as that obtained from the IV error correction approach. Note here that b_EY,IQ_ represents the error corrected *unstandardized regression effect* of 1 year of education on IQ. The variance in IQ accounted for by latent EY is 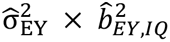 which must in turn be divided by the total variance in IQ in order to compute an R^2^ value. As the variance of a latent EduYears variable is not modelled in the IV approach, this R^2^ value cannot be directly computed in the IV approach.^1^

## Instrumental Variable Analysis for Error Correction of GPSs

DiPrete et al. (2017) propose using the same IV error correction approach described above to obtain an estimated of SNP heritability of EduYears that is disattenuated for measurement error by producing independent genome-wide polygenic scores (GPSs) for Educational Attainment (“multiple indicators”) in a prediction sample (HRS) computed from two sets of SNP regression weights, each computed from a different half of the EA2 meta-analytic sample (“by randomly splitting the GWAS sample that was used for [the GPS] construction”).

DiPrete et al. (2017) propose using one set of GPSs as the instrument for the second set of GPSs. Crucially, because the GPSs are calculated from different halves of the parent meta-analytic sample, their errors can be plausibly assumed to be uncorrelated, allowing one GPS to serve as an instrument for the other. This approach is represented in Figure 4, which is structurally equivalent to Figure 2. However, key here is that GPS scores do not have intrinsic units, and are therefore standardized with respect to their own sample distributions. For simplicity, EduYears are also standardized in this model, such that the coefficient can be interpreted in terms of SD units of EY. Error correction works as before, such that

**Figure 4.**
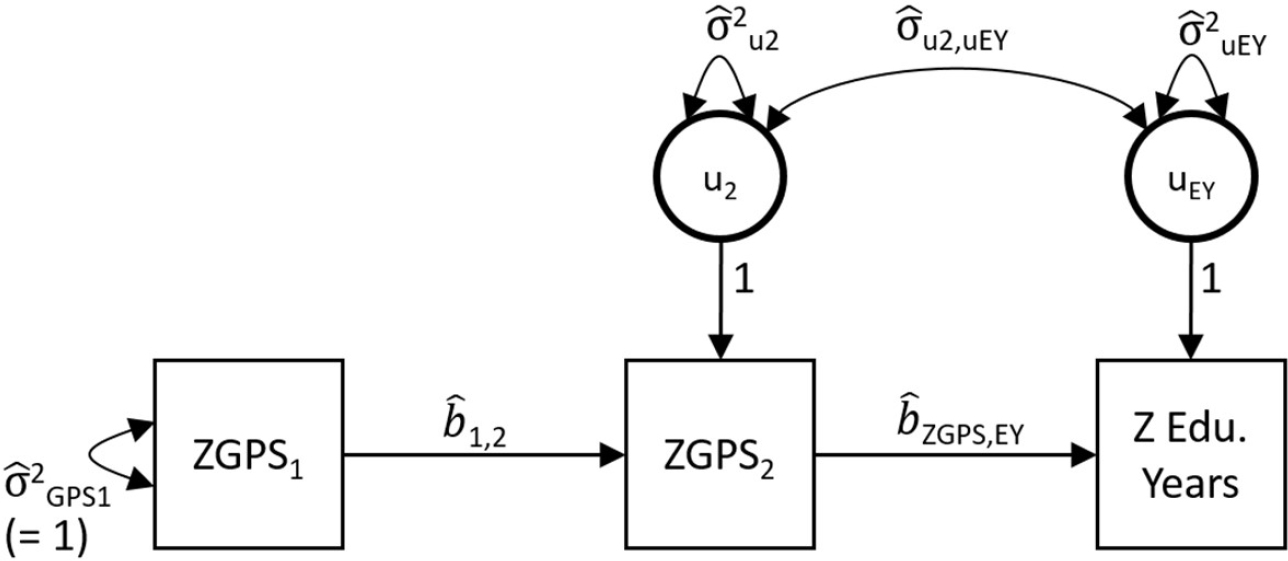
Note that all nonnumeric parameters in this figure are model estimates and are denoted by a hat (^). Numeric parameters are fixed values. Basic statistics that are calculated directly from the sample data have no hats.

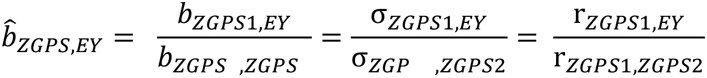

Key here is that 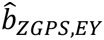 represents the error-corrected effect of one unit of ZGPS2 on ZEduYears. As ZGPS_2_ has been standardized with respect to the variance of ZGPS_2_, which is itself a mixture of true genetic variance and error variance, the regression effect has been corrected for measurement error but it is still in units of a score that has been standardized with respect to observed, as opposed to latent factor, variance. Thus the square of the 2SLS regression coefficient (i.e., 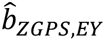) is not a correct estimate of the SNP heritability (see p. 9). Most generally, the square of a regression coefficient only represents R^2^ (i.e. SNP heritability) if it is a **standardized coefficient** from a univariate regression. ZEduYears has been appropriately standardized, but ZGPS is standardized with respect the variance of the observed indicator variable (GPS2), not the unobserved latent variable as it would need to be. As DiPret et al. (2017) explain, the corrected coefficient (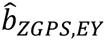) must both be squared and multiplied by an estimate of the variance of the unobserved latent genetic factor to rescale it to standardized units in order to obtain an appropriate estimate of the SNP heritability. This is made explicit in the psychometric parameterization presented next.

## An Explicit Psychometric Model for Error Correction of GPSs

As articulated earlier, in psychometrics multiple indicators are often leveraged as a means of error correction. In Figure 5, ZGPS1 and ZGPS2 are explicitly represented as imperfect indicators of a latent genetic factor, G, with uncorrelated errors. As in the earlier psychometric model, the metric of the latent variable (G) is linked to the metric of manifest variable EduYears1 via a fixed unstandardized loading of 1. If the two GPSs are computed from a randomly split discovery sample, we can make the simplifying (but unnecessary) assumption that the loadings are equivalent for ZGPS1 and ZGPS2. We can solve for the parameter 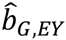 as follows:

**Figure 5.**
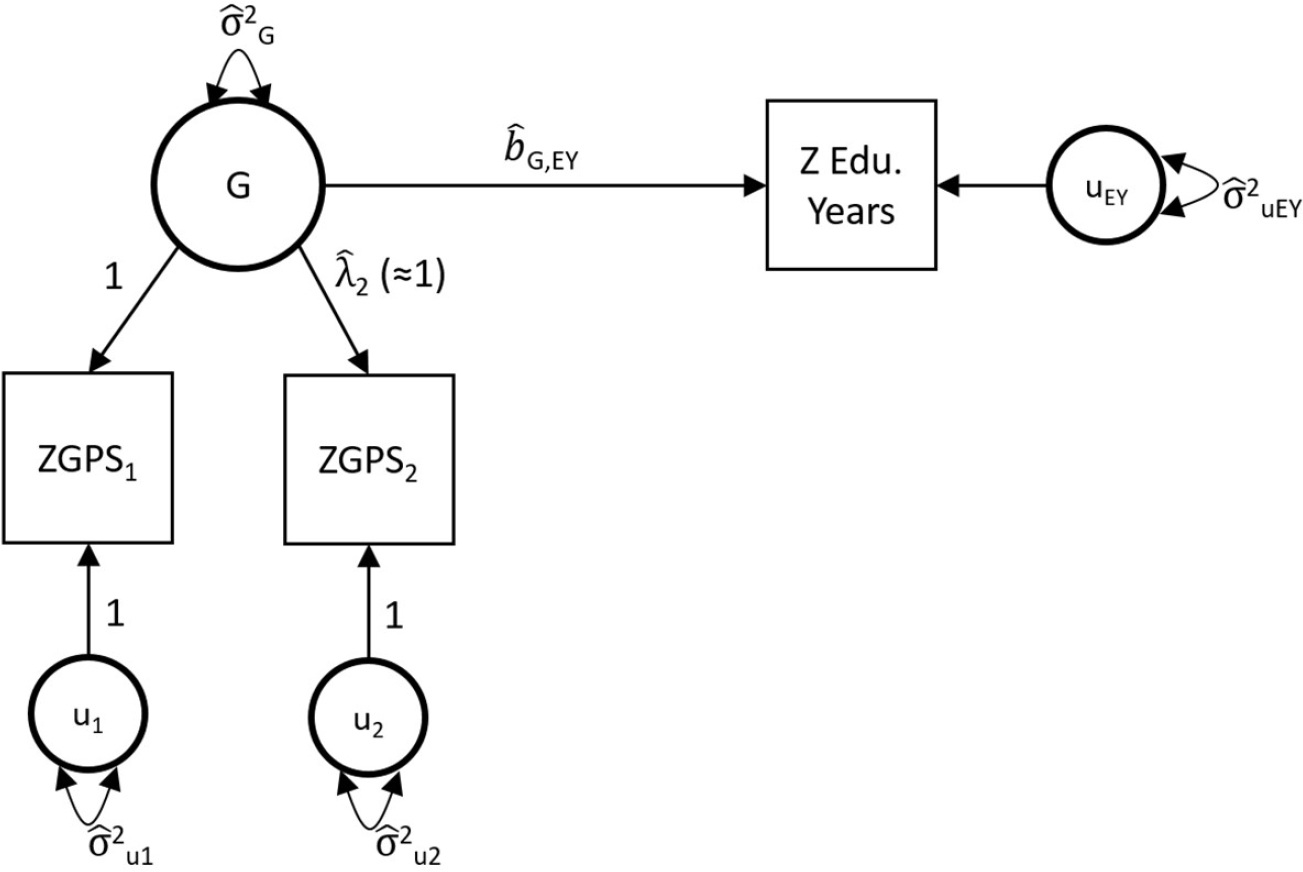
Note that all nonnumeric parameters in this figure are model estimates and are denoted by a hat (^). Numeric parameters are fixed values. Basic statistics that are calculated directly from the sample data have no hats.

**Figure 6.**
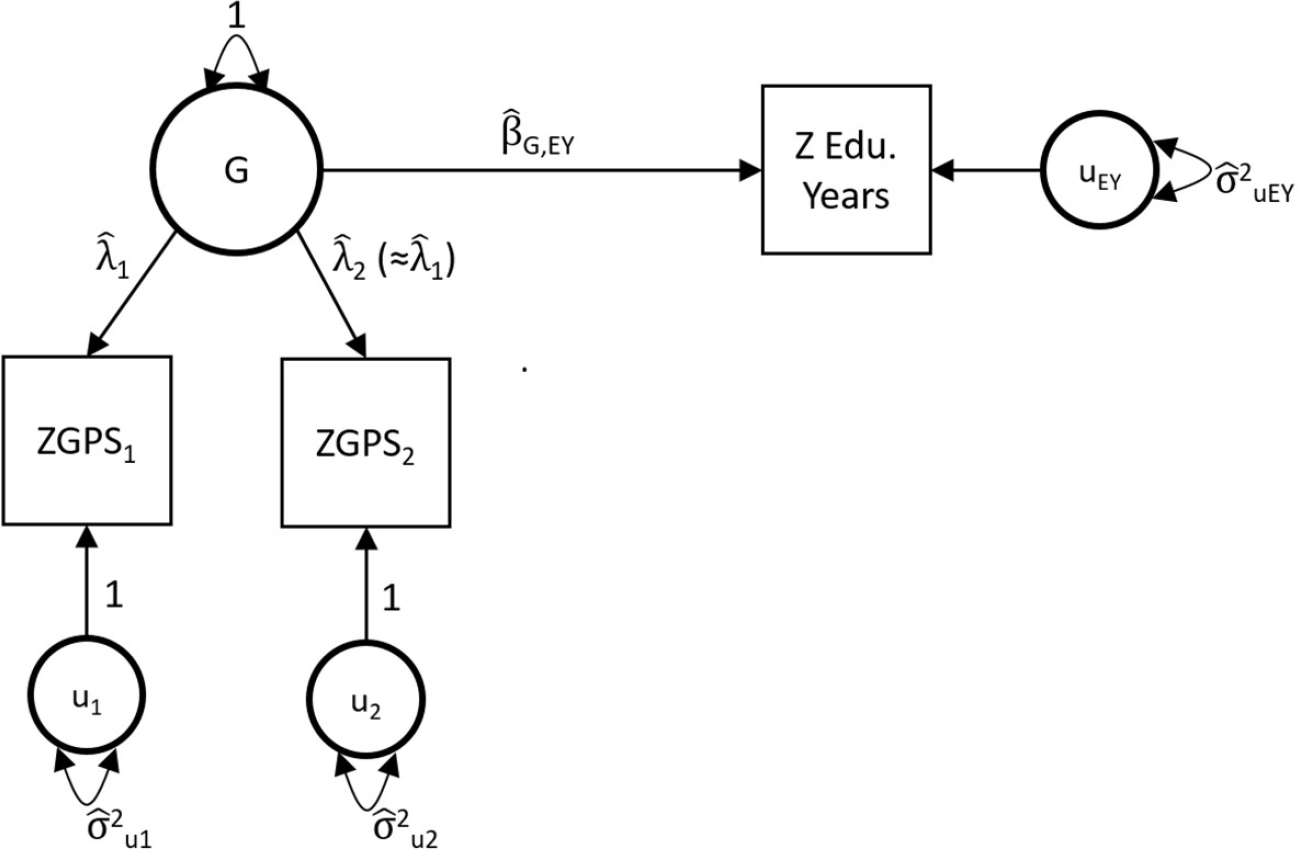
Note that all nonnumeric parameters in this figure are model estimates and are denoted by a hat (^). Numeric parameters are fixed values. Basic statistics that are calculated directly from the sample data have no hats.

**Figure 7.**
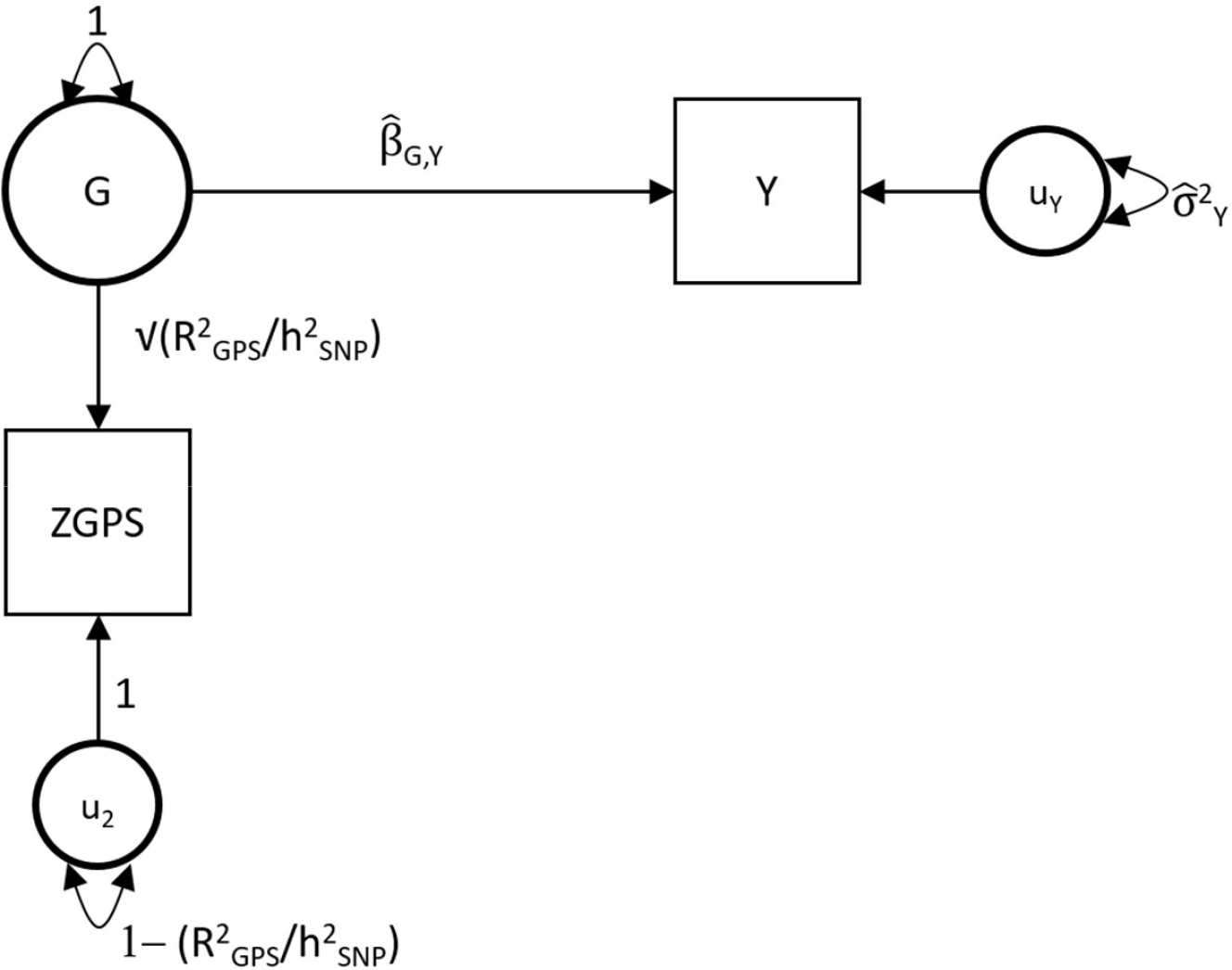
Note that all nonnumeric parameters with hats in this figure are model estimates and are denoted by a hat (^). The loading on G and the variance of u2 are fixed values that are computed outside of the model.

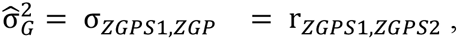

and

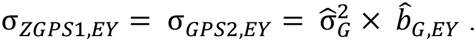

Thus

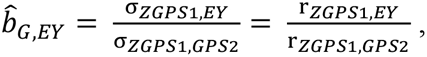

which is exactly the same as the corrected 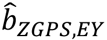 estimate obtained from the IV error correction approach.

Importantly 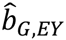 is an unstandardized estimate. To obtain an appropriate estimate of R^2^ (i.e. SNP heritability), the variance of the independent variable (the latent variable G) must be considered. As ZEduYears is already standardized, R^2^ is estimated as:

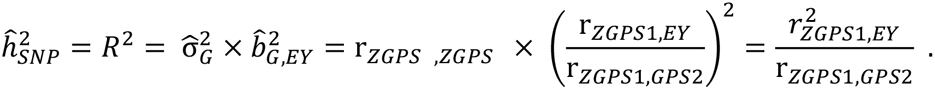

Note that the above equation, in which the estimate of 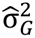 is equal to r_*ZGPS*1,*ZGPS*2_ only holds when the loadings of ZGPS1 and ZGPS2 are both equal to 1. If the discovery sample has been split by convenience, rather than randomly split, the loadings of ZGPS_1_ and ZGPS_2_ are less likely to be equivalent. This can easily be accommodated in the psychometric model presented in Figure 5, by allowing 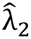 to be freely estimated.

## Respecifying the Psychometric Model to Standardized Units

The psychometric approach to measurement error correction can be easily re-specified in order to provide a more direct estimate of the standardized regression effect 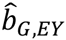 of G on the phenotype (i.e. the square root of the SNP heritability). Rather than defining the metric of G by linking it to the metric of ZGPS, we set its variance to 1, such that it is in standardized units. Again, we make the simplifying but unnecessary assumption that the loadings of ZGPS1 and ZGPS2 are equivalent. Under this specification:

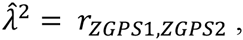

and

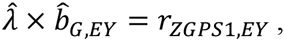

such that

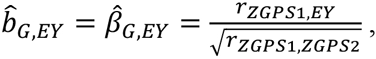

which squared produces the SNP heritability:

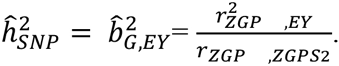

Interestingly, this result maps on directly to the well-known psychometric correction for measurement error attenuation (Spearman, 1904; https://en.wikipedia.org/wiki/Correction_for_attenuation), where r_ZGPS1,ZGPS2_ is a consistent estimate of the reliability of the GPSs, just as r_odd,even_ is a consistent estimate of the reliability of the two halves of a psychometric instrument using the split-half method (https://en.wikipedia.org/wiki/Reliability_(statistics). The attenuation correction formula allows for correction due to unreliability in both variables (i.e. GPS and EY), whereas the above formula only corrects for unreliability of the GPS (the reliability of EY is implicitly assumed to be 1.0, thus dropping out of the calculation):

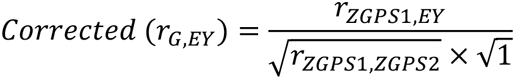

## Broader Application of the Psychometric Approach

The psychometric approach described above provides a framework for obtaining disattenuated estimates of associations between genetic risk, G, for the discovery phenotype (e.g. EduYears) and a novel phenotype, Y (e.g. Income). Thus, it can be leveraged for much more than obtaining an estimate of the SNP heritability of the discovery phenotype (for which an estimate can already be obtained using GREML or LD score regression in the parent meta-analytic sample). The novel phenotype is simply replaced as the dependent variable. The estimated regression effect of the latent genetic factor, G, on Y (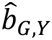) represents the disattenuated standardized effect of polygenic score for the discovery phenotype on the novel phenotype. Its square represents a disattenuated estimate of the proportion of variance explained in the novel phenotype (Income) by the polygenic score for the discovery phenotype (EduYears).

## Correcting GPSs for Error Using an External Estimate of SNP Heritability

Finally, the psychometric approach provides an instructive framework for conceptualizing how information about the SNP heritability of the discovery phenotype (EduYears) that is imported from LD Score regression or GREML in the parent meta-analytic sample (e.g. EA2) can be used to disattenuate a GPSs association with a novel dependent variable (e.g. income) in a prediction sample. Key here is that SNP heritability is assumed to be equivalent in magnitude across the parent discovery dataset and prediction dataset.

We no longer split the parent sample in half to estimate two GPSs and instead estimate a single GPS. We compute the reliability of the GPS as the ratio of the % of variance explained in the discovery phenotype in the prediction sample (e.g. the % of variance in EduYears explained by the GPS in HRS) to the SNP heritability estimated in the parent meta-analytic sample (e.g. the SNP heritability as estimated in EA2).

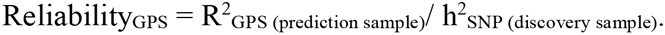

This estimate is imported into a structural equation model in the prediction dataset. ZGPS is specified to load on unit variance scaled latent factor (G) at a fixed value equal to the square root of Reliability_GPS_, and the residual variance in ZGPS is fixed to 1 - Reliability_GPS_. The estimated regression effect (b_G,Y_) of the latent genetic factor, G, on the novel phenotype Y represents the disattenuated standardized effect of polygenic score on the novel phenotype. Its square represents an estimate of the proportion of variance explained in the novel phenotype Y (e.g. Income) by the tagged SNPs relevant to the discovery phenotype (EduYears).

This model implies that

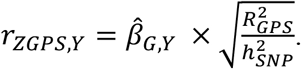

In other words, the observed association between the GPS and Y is attenuated by the square root of the GPS reliability. Solving for βG,Y yields:

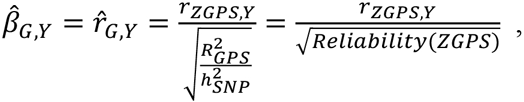

which is equivalent the well-known psychometric correction for measurement error attenuation in the independent variable (Spearman, 1904; https://en.wikipedia.org/wiki/Correction_for_attenuation).

Although not presented in this way, this is equivalent to what Daniel Benjamin, David Cesarini, and Patrick Turley (recently implemented by Beauchamp, 2016; *Online Supplement: “Directional Selection Differentials”*) have described as scaling of the regression coefficient by the SD of the true genetic score, rather than the SD of the observed GPS. This becomes clearer as follows:

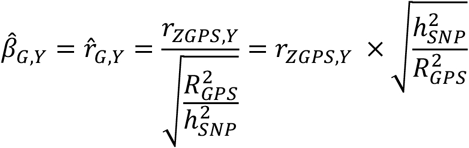

It is important to bear in mind that the both h^2^_SNP_ and R^2^_GPS_ are treated as fixed terms under this approach. However, each of these terms is actually estimated with error. Therefore, the standard error of disattenuated estimate 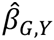 will be downwardly biased under this approach.

## A Conceptual Note

In order to be accurate, the above formulas to correct for disattenuation of GPS associations in predictions samples rely on the assumption that the individual SNP effects composing the GPS are uniform in their degree of error, or at least that the magnitude of imprecision of the individual SNP effects is unrelated to their effect sizes. I can envision circumstances in which this may not be the case. Consider, for example, a situation in which EduYears is affected by both heritable cognitive (e.g. IQ) and noncognitive factors (e.g. conscientiousness). Further, imagine a scenario in which the SNPs relevant to IQ have higher average minor allele frequencies (MAFs), and hence their coefficients have smaller standard errors, and the SNPs relevant to conscientiousness have smaller MAFs, and hence their coefficients have larger standard errors (cf. Penke & Jokela, 2016). Thus, the GPS for EduYears will more faithfully tag the SNPs relevant to IQ than it will for conscientiousness. We might be interested in then going on use the GPS for a different prediction phenotype than EduYears, such as IQ. We would calculate GPS-IQ association in a prediction sample and disattenuate it on the basis of reliability of the GPS. However, our correction may overestimate the association by compensating for the unreliability of the conscientiousness SNPs that were relevant for EduYears even though such SNPs are not as relevant for IQ. Daniel Benjamin has suggested that this limitation may be overcome by stratifying the correction by MAF bins.

## Footnotes

1 It is of further note that the IV approach requires choosing one of the two alternate forms (EY1 or EY2) as the IV. As this decision is arbitrary, it is prudent to run the IV approach both ways, and perhaps average the estimates. This is not necessary in the psychometric approach, although relaxation of the parallel alternative form assumption would allow for running the model twice, switching the anchor indicator.

